# Unveiling Gloriosine as a Dual-Acting Regulator of Glutamine Metabolism and Ferroptosis in Triple-Negative Breast Cancer: Insights from Network Pharmacology and Experimental Validation

**DOI:** 10.64898/2026.05.17.725321

**Authors:** Biswajit Dey, Essha Chatterjee, Bharat Goel, Shreyans Kumar Jain, Pradeep Kumar Naik, Santosh Kumar Guru

## Abstract

**Background:** TNBC lacks clearly defined molecular targets, so chemotherapy remains the standard approach despite resistance, toxicity, and high relapse rates. New, less toxic therapeutic options are urgently needed. This study evaluated the anticancer potential of gloriosine, a plant-derived alkaloid known to inhibit tumor cell growth with minimal effect on normal breast cells.

**Methods:** Putative targets of gloriosine were predicted using SwissTargetPrediction, TargetNet, and PharmMapper, then intersected with gene sets linked to TNBC and glutamine metabolism. The resulting network was characterized through protein-protein interaction mapping and Gene Ontology/KEGG enrichment analysis. Molecular docking assessed binding affinity to top targets, and predictions were validated experimentally using cell viability, colony formation, and wound-healing assays. Oxidative stress and ferroptosis were assessed via ROS (DCFDA), glutathione, and lipid peroxidation (MDA) assays, along with Western blotting and FerroOrange staining.

**Results:** Network analysis identified 100 predicted targets, 60 overlapping with TNBC/glutamine-metabolism genes; SRC, EGFR, mTOR, and HSP90AA1 emerged as hub proteins. Enrichment analysis linked this network to cancer progression, metabolic reprogramming, and resistance pathways, including central carbon metabolism and ErbB/EGFR-inhibitor resistance signaling. Docking confirmed strong gloriosine-target binding. Experimentally, gloriosine reduced cell proliferation and migration in a dose-dependent manner. Mechanistically, it suppressed glutaminolysis and induced ferroptosis, marked by increased ROS, glutathione depletion, elevated lipid peroxidation, GPX4 suppression, and intracellular iron accumulation.

**Conclusions:** Gloriosine suppresses TNBC growth via multi-target modulation and ferroptosis induction, supporting its potential as a novel anticancer candidate.

Graphical abstract
Flow chart of the network pharmacological and *in vitro* study of gloriosine

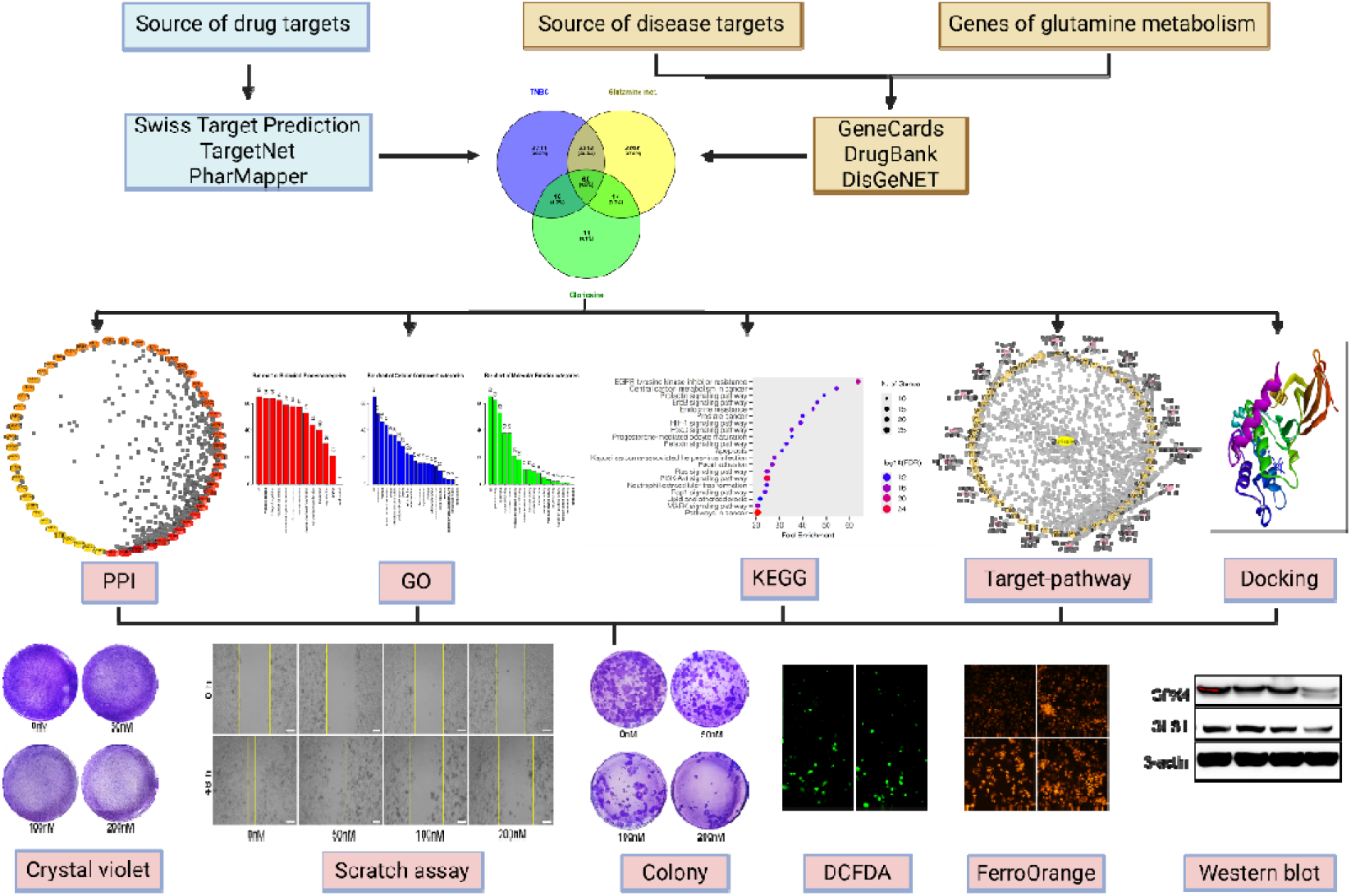

## Introduction

TNBC is an aggressive and heterogeneous subtype of breast cancer defined by the absence of estrogen receptor (ER), progesterone receptor (PR), and human epidermal growth factor receptor 2 (HER2) expression. This molecular profile significantly limits the applicability of targeted therapies, rendering cytotoxic chemotherapy the mainstay of systemic treatment. Despite initial responses, TNBC is frequently associated with the rapid development of chemoresistance, high rates of recurrence, and early metastasis, highlighting the urgent need for novel therapeutic agents with distinct and more effective mechanisms of action [1][2].

Natural products have long been a valuable source of anticancer agents, offering structural diversity and multi-targeted therapeutic potential. Gloriosine, a bioactive alkaloid isolated from *Gloriosa superba*, has recently gained attention due to its potent antimitotic and cytotoxic properties. Gloriosine, structurally similar to colchicine, binds to tubulin and interferes with microtubule dynamics, leading to impaired mitotic spindle formation and cell cycle arrest at metaphase [3][4]. This mode of action is particularly relevant in TNBC, where rapid cellular proliferation necessitates efficient mitotic progression. Furthermore, natural compounds such as gloriosine often exert pleiotropic effects, enabling them to modulate multiple signaling pathways simultaneously and potentially overcome resistance mechanisms that compromise the efficacy of conventional chemotherapeutic agents [5][6].

In addition to uncontrolled proliferation, metabolic reprogramming is a hallmark of TNBC. Among these alterations, glutamine metabolism plays a critical role in supporting tumor growth and survival by fueling the tricarboxylic acid (TCA) cycle, maintaining redox balance, and promoting biosynthetic processes [7]. TNBC cells often exhibit increased glutamine dependency, making glutaminolysis a potential therapeutic vulnerability. Targeting key enzymes involved in glutamine metabolism, such as glutaminase and glutamate dehydrogenase, has been shown to impair tumor progression and sensitize cancer cells to treatment [8]. Furthermore, dysregulated glutamine metabolism contributes to redox homeostasis, thereby influencing susceptibility to oxidative stress-mediated cell death pathways [9].

Ferroptosis is an iron-dependent mode of cell death characterized by the excess accumulation of lipid peroxides. [10]. It is characterized by oxidative damage to cellular membranes and has been shown to be particularly relevant in TNBC cells. TNBC cells are especially susceptible to ferroptosis due to their altered redox homeostasis and elevated iron metabolism [11]. Notably, glutamine metabolism has been reported to facilitate ferroptosis by contributing to mitochondrial activity and reactive oxygen species (ROS) generation [12], further linking metabolic reprogramming with cell death regulation. Inducing ferroptosis could therefore represent a promising strategy to eliminate TNBC populations. Compounds capable of simultaneously disrupting mitosis, altering metabolic pathways, and promoting oxidative stress may exert synergistic anticancer effects.

Advancements in computational biology have further accelerated drug discovery through systems-level approaches. Network pharmacology, which integrates pharmacology with systems biology, artificial intelligence, and big data analytics, enables the identification of complex interactions between drugs, targets, and diseases [13][14]. By constructing and analyzing “drug–target–disease” networks, this approach provides valuable insights into the molecular mechanisms underlying the therapeutic effects of bioactive compounds, particularly those derived from natural sources. Such integrative analyses are instrumental in uncovering key signaling pathways and potential biomarkers involved in disease modulation.

In this context, the present study seeks to systematically evaluate the therapeutic potential of gloriosine against TNBC using an integrated approach of network pharmacology, molecular docking, and in vitro experimental validation. Through this integrated approach, we seek to identify the key molecular targets and signaling pathways associated with gloriosine, to develop this compound as a novel chemotherapeutic agent for TNBC therapy.

## Materials and methods

### Source of drug targets

The SwissTargetPrediction database (http://www.swisstargetprediction.ch/) [15], TargetNet (http://targetnet.scbdd.com/) [16] and PharmMapper database (http://www.lilab-ecust.cn/pharmmapper/) [17] were employed to identify potential targets of gloriosine, using a probability threshold greater than 0 as the screening criterion. The UniProt database (https://www.uniprot.org/) [18] was subsequently utilized to standardize and map the gene names of the predicted target proteins. Finally, the targets identified from all three databases were combined to compile a comprehensive list of gloriosine-associated drug targets.

### Sources of disease targets

The databases GeneCards (https://www.genecards.org/) [19] DrugBank (https://go.drugbank.com/) [20] and DisGeNET (https://www.disgenet.org/) [21], were explored to identify relevant disease-associated targets. Using the standard search options in each database, all TNBC and metabolism-related targets were collected. The results from these three sources were then integrated to compile a unified set of disease-associated targets. Subsequently, these disease targets and metabolic genes were intersected with the identified drug targets to determine the overlapping targets. A Venn diagram illustrating the intersection between drug, disease and glutamine metabolism genes was generated using Venny 2.1.0 (https://bioinfogp.cnb.csic.es/tools/venny/).

### Construction of PPI (Protein-Protein Interaction) network

A total 60 genes that showed the relevance score between the drug and the disease were uploaded to the STRING database (https://string-db.org/) [22], with the organism parameter set to *Homo sapiens* and the confidence score threshold adjusted to more than 0.4. Isolated nodes were removed, and a PPI (protein-protein interaction) network was generated. The resulting interaction data were then imported into Cytoscape 3.8.0 for topological analysis, where node degree values were calculated, and the PPI network was visualized among these genes.

### GO (Gene Ontology) functional enrichment analysis

The intersecting targets between the drug and disease were uploaded into the WebGestalt database (http://www.webgestalt.org/) [23]. The analysis was limited to Homo sapiens as the selected species. Over-representation analysis (ORA) was employed as the analytical method. Using the genomic protein-coding genes as the reference set, Gene Ontology (GO) enrichment analysis was performed to evaluate the associated biological processes (BP), molecular functions (MF), and cellular components (CC) of the identified intersection targets.

### KEGG (Kyoto Encyclopedia of Genes and Genomes) pathway enrichment analysis

The intersecting targets were imported into the ShinyGO online tool (https://bioinformatics.sdstate.edu/go/) [24] with the species restricted to *Homo sapiens*. Significance criteria were set at p < 0.05 and a false discovery rate (FDR) < 0.05. KEGG pathway enrichment analysis was then performed on these targets.

### Molecular docking of drug components intersecting targets

Molecular docking studies were performed using AutoDock 4.2 to understand the molecular interaction between the ligand (gloriosine) and proteins, viz. EGFR (PDB: 2J6M), mTOR (PDB: 4JT6), SRC (PDB: 2BDJ), and HSP90α (PDB: 3O0I) [25]. The ligand molecule was sketched in ChemDraw, followed by energy minimization using the MM2 force field, and then saved in MOL2 format. The crystal structures of proteins were obtained from the Protein Data Bank (https://www.rcsb.org/). All the water molecules, ligands, and ions were removed. Non-polar hydrogen atoms were removed, and Gasteiger charges were added using M.G.L Tools 1.5.7. Blind docking was performed to explore all potential binding sites on the receptor. The grid box was centered at the geometric center of the protein and expanded to dimensions of 120 × 120 × 120 grid points, fully enclosing the entire macromolecule structure. The grid point spacing was maintained at the default value of 0.375 Å. The docking study was performed using Lamarckian Genetic Algorithm (LGA). The docking was performed with 100 runs, 150 population size, 27,000 number of generations, and 2,500,000 number of energy evaluation [26]. It employs a ‘semiempirical free energy force field’ to evaluate conformations at the time of docking simulation. The docked pose was visualized by Discovery Studio Visualizer for studying interactions.

### Cell lines and cell culture

The human TNBC cell lines MDA-MB-231 (HTB-26, RRID: CVCL_0062) and Hs 578T (HTB-126, RRID: CVCL_0332) were sourced from the American Type Culture Collection (ATCC). MDA-MB-231 cells were maintained in RPMI 1640 medium (Gibco, cat. 11875093) supplemented with 10% heat-inactivated fetal bovine serum (FBS; Gibco, cat. 26140079) and penicillin/streptomycin (ThermoFisher Scientific, cat. 15140122). Additionally, 2 mM glutamine (cat. 25030081, Gibco, ThermoFisher Scientific) and 2 g/L sodium bicarbonate (Sigma, cat. S5761) were included. All cultures were grown at 37 °C in a humidified incubator with 5% CO□. Cells were cultured and treated with gloriosine alone or in combination with BPTES (TargetMol, cat. T6791), ferrostatin-1 (TargetMol, cat. T6500) and erastin (TargetMol, cat. T1765) for the experiments.

### Cell viability assay

MTT assay was performed to determine the cell viability. Briefly, cells were seeded in 96-well plates, and after the 24 h incubation, were treated with different concentrations of gloriosine. Following 48 h treatment, MTT reagent (2.5 mg/mL) was added for 3–4 h at 37°C. The resulting formazan crystals were dissolved in DMSO, and absorbance was measured at 570 nm using a microplate reader (EnVision) to determine relative cell viability.

### Wound healing assay

The wound-healing migration assay was conducted as previously described [27]. 6-well plates were used to seed cells. Once the cells reached approximately 90% confluence, a scratch was introduced using a 200 µL pipette tip. PBS was used to wash the wells, following treatment. Images were acquired after 48 h of treatment using a microscope (AxioVert A1).

### Colony formation assay

Cells were treated with different concentrations of gloriosine for 48 hours. After treatment, cells were trypsinized, and about 500 cells were replated into 6-well plates and incubated for two weeks to allow colony growth. The resulting colonies were then fixed using 25% methanol, stained with 0.5% crystal violet, and those consisting of 50 or more cells were counted across three independent experiments [28].

### Determination of GSH levels

Intracellular levels of reduced glutathione (GSH) were measured using Ellman’s reagent [5,5′-dithiobis-(2-nitrobenzoic acid), DTNB] [29]. In brief, cells were harvested, washed with PBS, and lysed in ice-cold RIPA buffer. The lysates were then deproteinized using 5% sulfosalicylic acid and centrifuged at 10,000×g for 10 min at 4 °C. The resulting supernatants were collected for analysis. For the assay, sample aliquots or GSH standards were incubated with DTNB (final concentration approximately 0.5 mM) in phosphate buffer. The production of the yellow-colored 5-thio-2-nitrobenzoic acid (TNB) was measured by recording absorbance at 412 nm using a microplate reader. GSH levels were determined from a standard curve generated with known concentrations of reduced GSH and were normalized to total protein content as measured by the Bradford assay.

### Estimation of lipid peroxidation

Lipid peroxidation levels were determined by measuring malondialdehyde (MDA) formation as described previously with minor modifications [30]. Briefly, 50 μL of cell lysate was mixed with a solution containing 0.67% thiobarbituric acid (TBA) and 20% trichloroacetic acid (TCA), followed by incubation at 95°C for 40 min. After cooling, the absorbance of the reaction mixture was recorded at 532 nm. MDA concentrations were calculated using a standard curve generated from 1,1,3,3-tetramethoxypropane (TMP). Protein concentrations were determined using the Bradford assay, and the results were expressed as nanomoles of MDA per milligram of protein.

### FerroOrange staining

Cells were seeded in 96-well plates and allowed to adhere for 24 h. Following this incubation period, the cells were treated with the compound for an additional 48 h. After treatment, cells were co-incubated with FerroOrange (1 mM) at 37 °C for 30 min. Subsequently, the cells were washed three times with PBS to remove excess dye, and intracellular Fe²□ levels were assessed by quantifying fluorescence signals using a HCS imaging system (cat. HCSDCX7LZRPRO) [31].

### Preparation of cell lysates and Western blot analysis

Cells were cultured in 100 mm petri dishes and treated at approximately 60% confluency with gloriosine for 48 h. After treatment, cells were washed with PBS, trypsinized, and harvested for lysate preparation. Protein extracts were prepared using RIPA buffer (cat. R0278, Sigma), and concentrations were quantified using the Bradford assay. The lysates were then mixed with loading buffer and heated at 95 °C for 10 min Equal amounts of protein (60 μg) were resolved by SDS-PAGE and subsequently transferred onto PVDF membranes. Membranes were blocked with 3% BSA in TBST and incubated overnight at 4 °C with the appropriate primary antibody (GPX4, cat. 52455S. GLS1, cat. 49363S. β-actin, cat. A5441). Following washes, membranes were incubated with HRP-conjugated secondary antibody (cat. 7074S), and protein bands were detected using a ChemiDoc imaging system (Bio-Rad) [28].

### Gene expression analysis by qRT-PCR

MDA-MB-231 and HCC1395 breast cancer cells were exposed to gloriosine for 48 h prior to RNA isolation. Total RNA was extracted using TRI Reagent (Sigma, Cat. No. T9424) according to the manufacturer’s protocol. First-strand cDNA was synthesized from the isolated RNA using the RevertAid cDNA Synthesis Kit following the supplier’s instructions. Quantitative real-time PCR (qRT-PCR) was subsequently carried out using SYBR Green Master Mix on a QuantStudio 7 Pro (Thermo Fisher Scientific). The primers SLC7A11 (forward 5’-TCTCCAAAGGAGGTTACCTGC-3’, reverse 5’-AGACTCCCCTCAGTAAAGTGAC-3’) and GPX4 (forward 5’-GAGGCAAGACCGAAGTAAACTAC-3’, reverse 5’-CCGAACTGGTTACACGGGAA-3’) were used in this study, and gene expression levels were normalized to GAPDH (forward 5’-GGAGCGAGATCCCTCCAAAAT-3’, reverse 5’-GGCTGTTGTCATACTTCTCATGG-3’) as the internal reference control.

### Metabolite detection assay

Intracellular glutamine and glutamate levels were quantified using the Glutamine/Glutamate-Glo Assay (Promega, cat. J8021) according to the manufacturer’s instructions. Cell lysates were prepared after the treatment and incubated with glutaminase + glutaminase buffer or glutaminase buffer alone, followed by the addition of the Glutamate Detection Reagent. After incubation at room temperature, luminescence was measured using a microplate luminometer. Glutamine concentrations were calculated by subtracting glutamate values from the total glutamine plus glutamate signal, and metabolite levels were normalized to protein content.

### Graphical diagram making

All graphical diagrams were made using Biorender (https://www.biorender.com/).

### Statistical analysis

Data are expressed as mean ± standard deviation (SD). The Dunnett method was employed for the one-way ANOVA test using GraphPad 8. A p-value of <0.05 was considered significant (*p□<□0.05; **p□<□0.01; ***p□<□0.001).

## Results

### Drug targets

Using the phytochemical structure of gloriosine as input, potential protein targets were predicted using SwissTargetPrediction, TargetNet and PharmMapper databases. This initial analysis identified a total of 100 candidate targets. To ensure consistency and accuracy, all predicted protein targets were subsequently standardized using the UniProt database. This normalization step removed redundancies and resolved inconsistencies in gene and protein nomenclature across the datasets, resulting in a refined and reliable target list.

### Identification of disease-associated and glutamine metabolism targets and their intersection with drug targets

A total of 4,698 TNBC-associated targets and 5,252 glutamine metabolism-related targets were retrieved from the GeneCards, DrugBank, and DisGeNET databases. Following data integration, these targets were intersected with the predicted gloriosine-associated targets. This analysis identified 60 overlapping targets, which are considered potential therapeutic targets of gloriosine in the treatment of TNBC (Figure 1).

**Figure 1:**
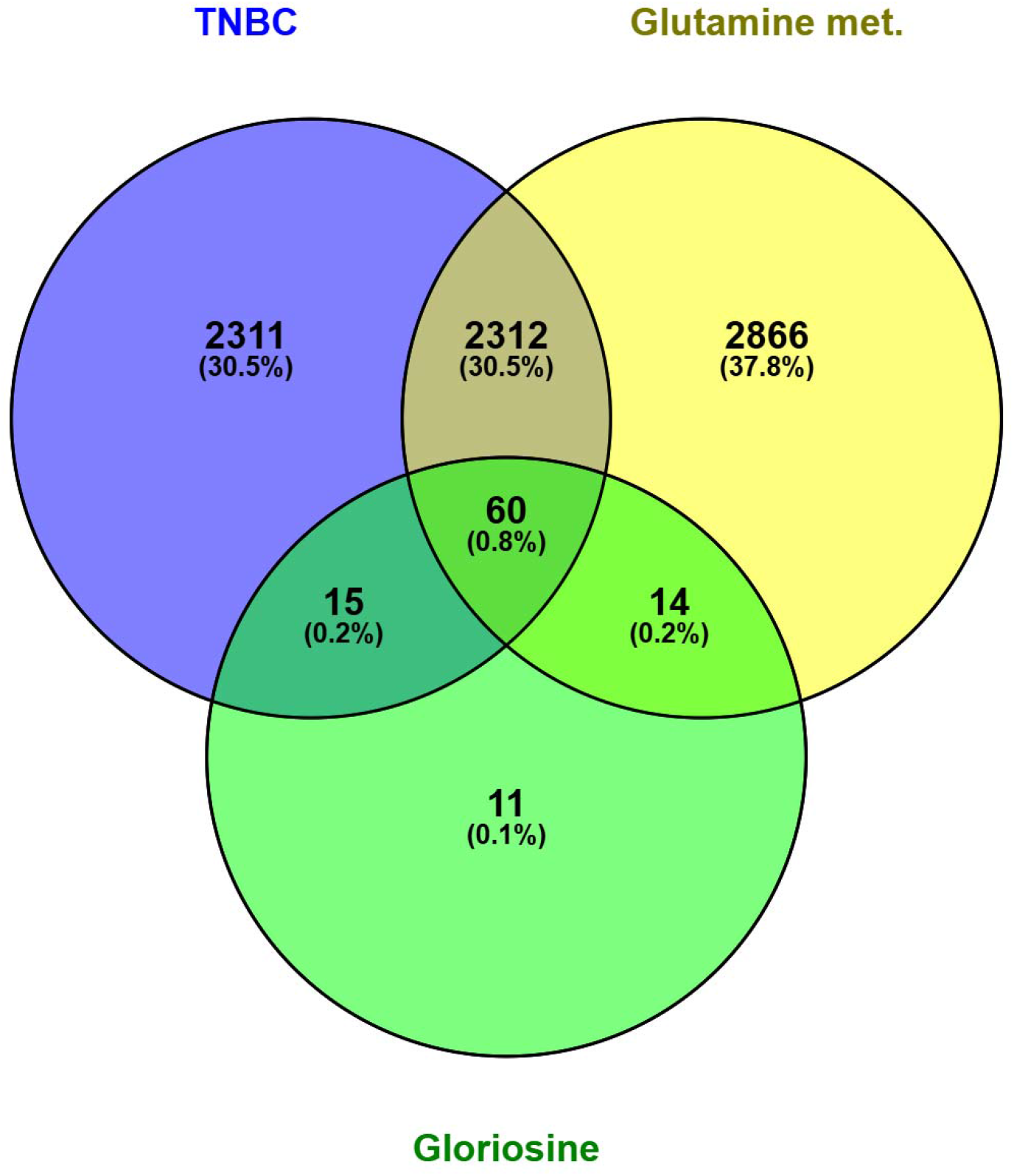
Venn diagram illustrating the overlapping targets among gloriosine, glutamine metabolism–associated genes, and triple-negative breast cancer (TNBC)

### Construction and analysis of the PPI network

The 60 overlapping targets were uploaded to the STRING database to construct a protein-protein interaction (PPI) network. The resulting network comprised 60 nodes, representing the potential interactions among gloriosine-associated targets in TNBC, as illustrated in Figure 2A. The network data were subsequently exported to Cytoscape 3.8.0 for topological analysis. Degree values were calculated to evaluate the relative importance of each node within the network. The results indicated that targets such as PARP3 (Poly(ADP-ribose) Polymerase 3), ABCC1 (ATP Binding Cassette Subfamily C Member 1), TGM2 (Transglutaminase 2), PGK1 (Phosphoglycerate Kinase 1), PDE4D (Phosphodiesterase 4D), Calpain 1 (CAPN1), P2RX7 (Purinergic Receptor P2X 7), GRM1 (Glutamate Metabotropic Receptor 1), NR3C2 (Nuclear Receptor Subfamily 3 Group C Member 2), and CTSL (Cathepsin L) exhibited low connectivity, with degree values ≤ 5. In contrast, targets including FPR1 (Formyl Peptide Receptor 1), CYP19A1 (Cytochrome P450 Family 19 Subfamily A Member 1), MMP1 (Matrix Metallopeptidase 1), LRRK2 (Leucine-Rich Repeat Kinase 2), MAPK9 (Mitogen-Activated Protein Kinase 9), FLT4 (Fms Related Receptor Tyrosine Kinase 4), NOS2 (Nitric Oxide Synthase 2), and ADAM17 (ADAM Metallopeptidase Domain 17) showed moderate connectivity, with degree values ranging between 5 and 9.9. The remaining targets demonstrated high connectivity, with degree values greater than 10. Notably, the highest degree values (≥ 40) were observed for SRC (SRC Proto-Oncogene, Non-Receptor Tyrosine Kinase), EGFR (Epidermal Growth Factor Receptor), mTOR (Mechanistic Target of Rapamycin), and HSP90AA1 (Heat Shock Protein 90 Alpha Family Class A Member 1), suggesting their central roles in the PPI network (Figure 2B).

**Figure 2:**
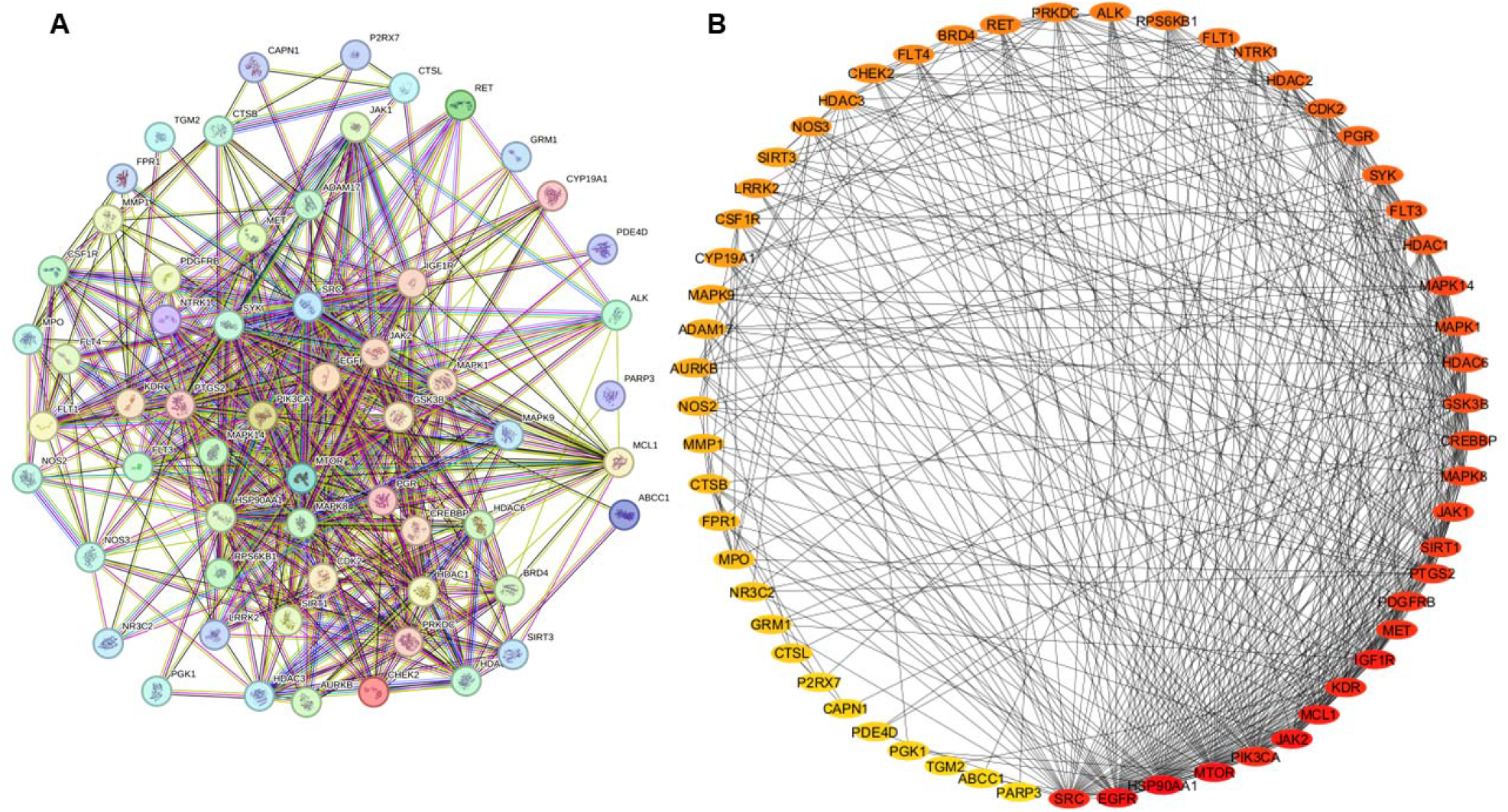
Protein–protein interaction (PPI) network of intersecting targets of gloriosine in TNBC associated with glutamine metabolism. **(A)** PPI network constructed using the STRING database. **(B)** network visualization in Cytoscape 3.10.3, where node color yellow to red represents increasing degree of connectivity.

### GO functional enrichment analysis

The 60 overlapping targets were analyzed using the WebGestalt database for GO functional enrichment, and the results are presented in Figure 3A. A total of 12 BP were identified for gloriosine in TNBC treatment, primarily related to metabolic processes, biological regulation, response to stimuli, multicellular organismal processes, and development. Additionally, 20 CC were enriched, mainly associated with the nucleus, cell membrane, protein-containing complexes, cytoplasm, membrane lumen, and endomembrane system. Furthermore, 17 MF were identified, largely involving protein binding, ion binding, nucleotide binding, transferase activity, and molecular transducer activity.

**Figure 3:**
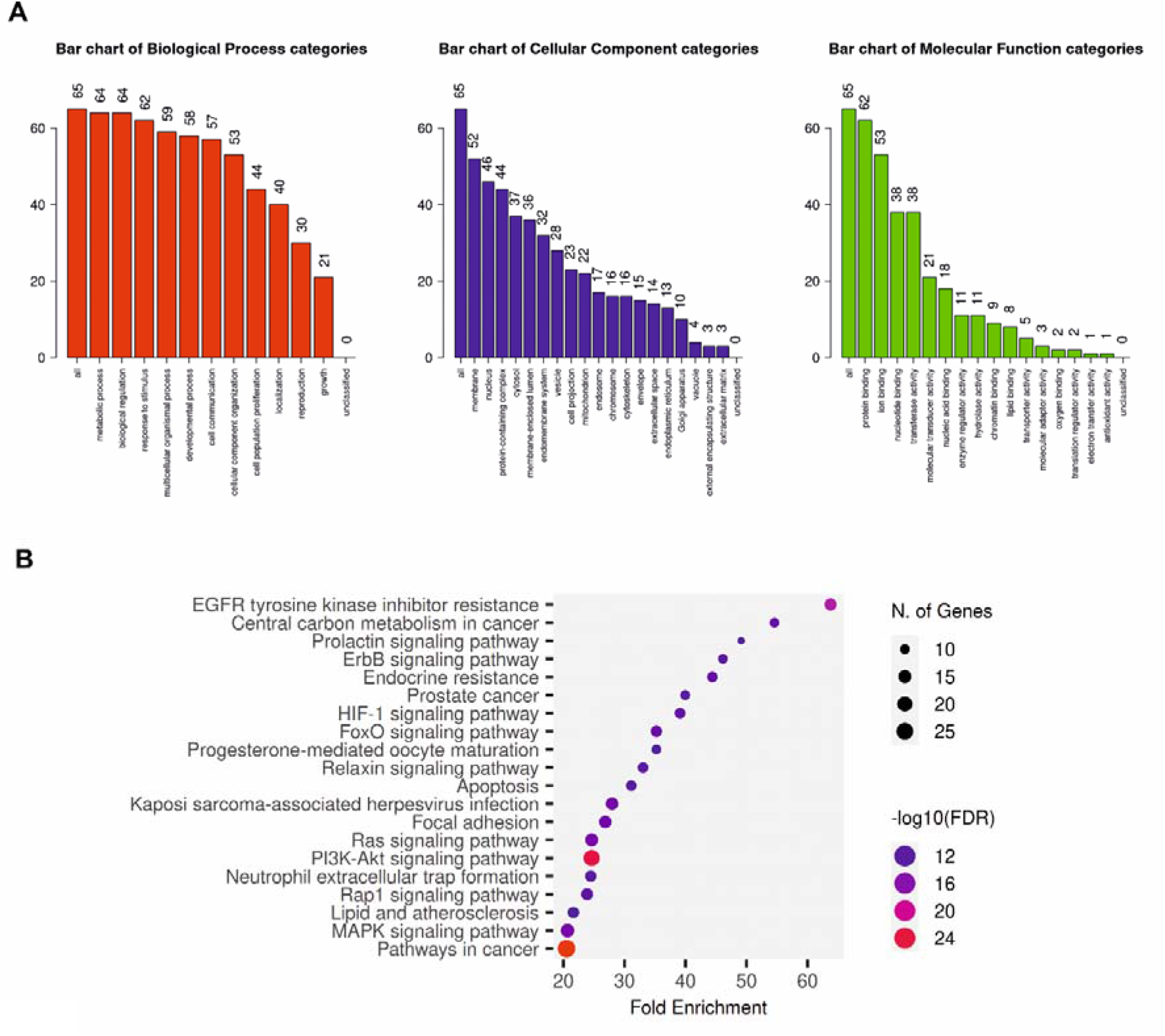
GO and KEGG pathway analysis. **(A)** Gene Ontology (GO) enrichment analysis of gloriosine-targeted genes associated with glutamine metabolism in triple-negative breast cancer (TNBC), showing the top enriched functional categories for biological processes (BP, red), cellular components (CC, blue), and molecular functions (MF, green). **(B)** KEGG pathway enrichment analysis of gloriosine-targeted genes involved in glutamine metabolism in TNBC, highlighting the significantly enriched signaling pathways associated with cancer progression and metabolic regulation.

### KEGG pathway enrichment analysis

The 60 overlapping targets were analyzed using the ShinyGO gene set enrichment tool for KEGG pathway enrichment. The top 20 pathways associated with gloriosine in TNBC treatment are presented in Figure 3B, including EGFR tyrosine kinase inhibitor resistance, central carbon metabolism in cancer, prolactin signaling, ErbB signaling, and endocrine resistance. The integrated network of drug-target-pathway interactions is illustrated in Figure 4.

**Figure 4:**
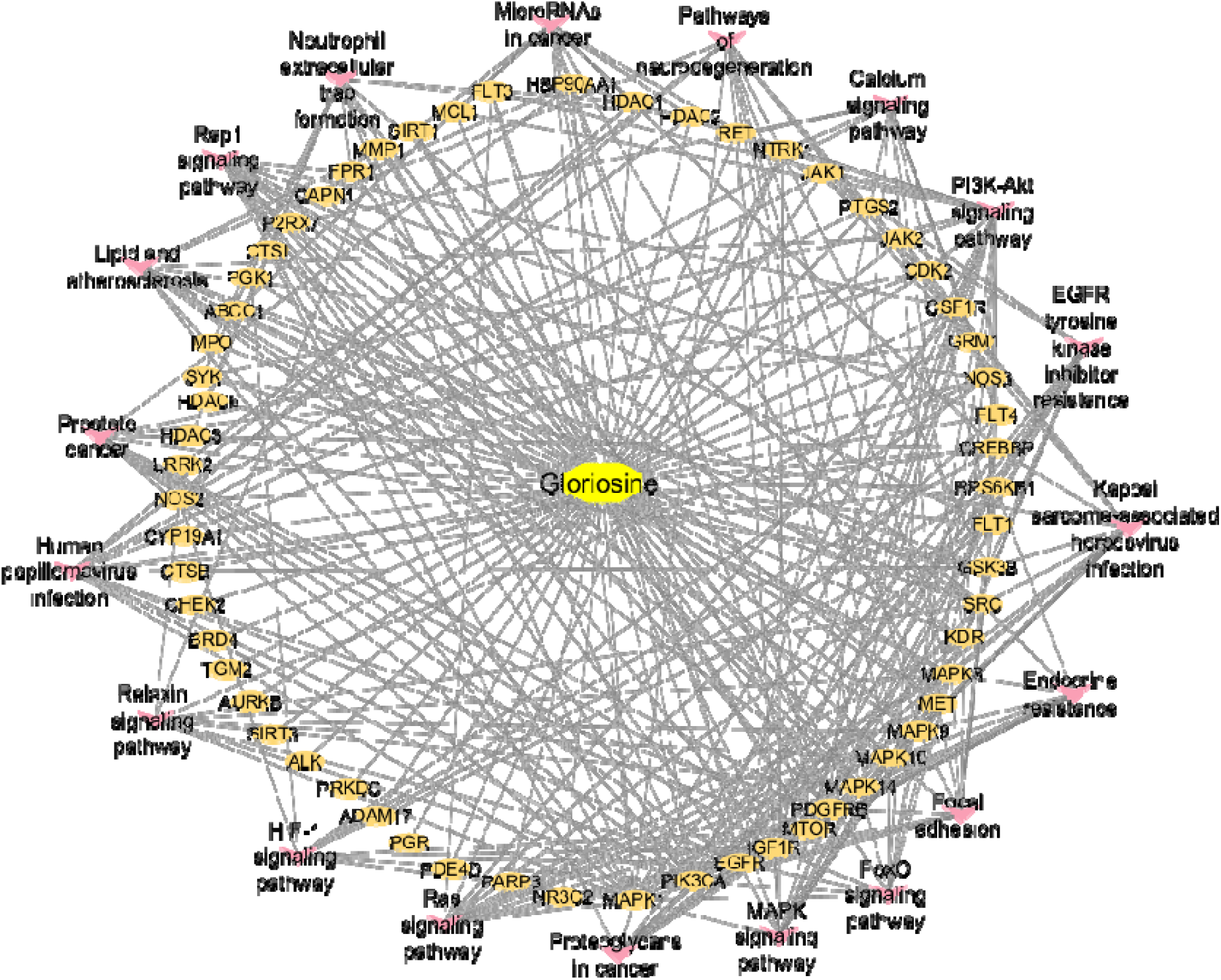
The network illustrates interactions between gloriosine, glutamine metabolism regulatory genes and associated signaling pathways in TNBC. Nodes represent the drug, targets, and pathways, while edges indicate their interactions, highlighting the multi-target role of gloriosine in modulating glutamine metabolism.

### Molecular docking analysis to validate binding affinity

Molecular docking analysis was carried out to evaluate the binding affinity of gloriosine against those targets which has highest degree value (EGFR, mTOR, SRC, and HSP90AA1). The docking results revealed that gloriosine exhibited favorable binding interactions with all selected targets, with binding energies ranging from −6.73 to −9.18 kcal/mol. Among the studied proteins, the strongest binding affinity was observed with HSP90AA1 (PDB ID: 3O0I), which showed a binding energy of −9.18 kcal/mol, indicating a highly stable ligand–protein complex (Figure 5A). Gloriosine also showed significant binding affinity toward mTOR (PDB ID: 4JT6), exhibiting binding energies of −9.01 kcal/mol (Figure 5B). Moderate binding affinities were observed with SRC (PDB: 2BDJ) and EGFR (PDB: 2J6M) with binding energies of −7.93, and −6.73 kcal/mol, respectively (Figures 5C and 5D). The ligand showed important interactions with the proteins viz. H-bond, van der Waals, π-alkyl, π-π-stacked, and π-cation. Overall, the docking results suggest that gloriosine is capable of interacting with multiple protein targets, showing a broad spectrum of binding affinities.

**Figure 5:**
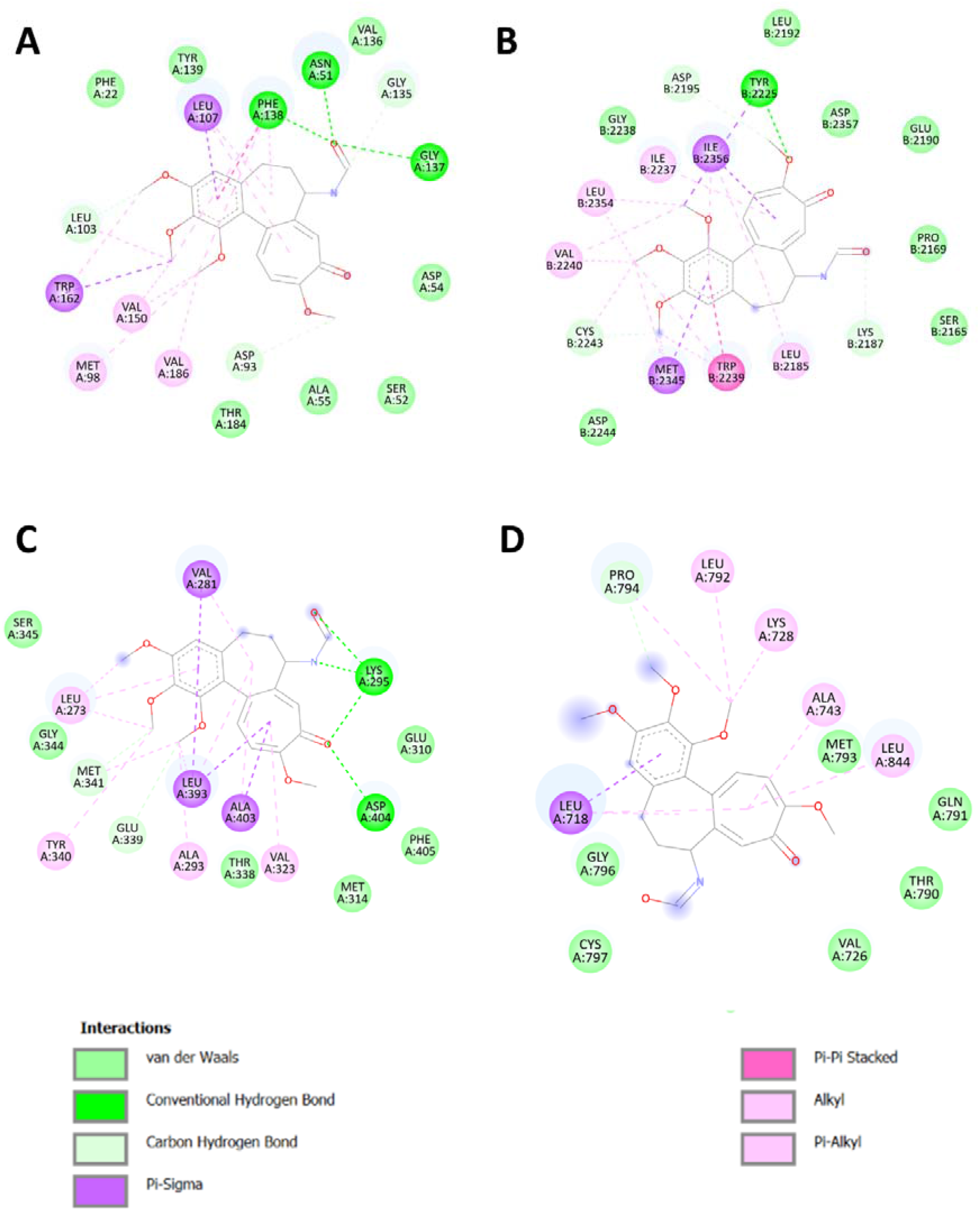
Molecular docking analysis of gloriosine with target proteins. Two-dimensional (2D) and three-dimensional (3D) interaction profiles illustrating the binding of gloriosine with key target proteins: **(A)** HSP90AA1, **(B)** mTOR, **(C)** SRC, and **(D)** EGFR.

### Gloriosine inhibits the proliferation and migration of TNBC cells

Gloriosine demonstrated inhibitory effects on cell proliferation different TNBC cell lines, while exhibiting a comparatively higher IC□□ value in normal breast epithelial MCF 10A cells (Table 1). Specifically, gloriosine displayed an IC□□ of approximately 100 nM in MDA-MB-231, 88 nM in HCC1395 and 98 nM in Hs 578T cells, indicating its potent cytotoxic effect on TNBC cell lines. To evaluate the effect of gloriosine on cell migration, a wound healing assay was performed. The results revealed a concentration-dependent inhibition of migratory ability in MDA-MB-231 and HCC1395 cells following treatment with gloriosine (Figure 6A). Colony formation assay further supported the concentration-dependent inhibition of cell proliferation upon gloriosine treatment in both cells (Figure 6B). The overall findings represented in Figure 6C.

**Figure 6:**
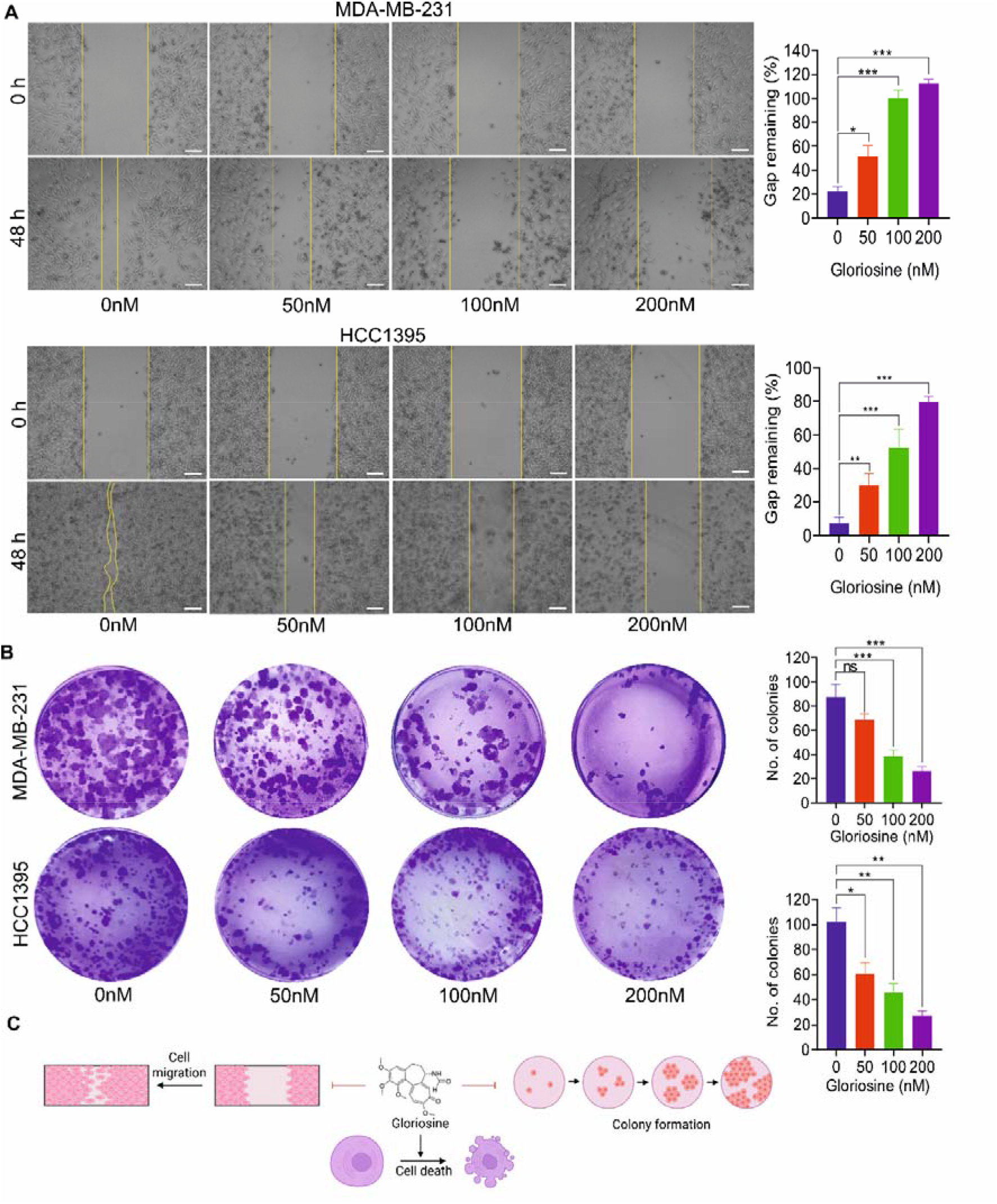
Gloriosine inhibits cell proliferation and migration. **(A)** Gloriosine inhibits cell migration in a dose-dependent manner. Scale bar = 50 µm. **(B)** Colony formation was hindered by the compound in a concentration-dependent manner. Data represent the Mean ± SD on three independent experiments (one-way ANOVA, *P<0.05, **P□<□0.01, ***P□<□0.001). **(C)** Summary represented in Fig. 6.

**Table 1:** Cytotoxic activity of gloriosine across different cancer cell lines and normal breast epithelial cells. The half-maximal inhibitory concentration (IC□□) values of gloriosine were determined in different cancer cell lines, along with the normal breast epithelial cell line MCF 10A. Cells were treated with increasing concentrations of gloriosine, and cell viability was assessed after the indicated treatment period using the MTT assay. IC□□ values are expressed in nanomolar (nM) as mean ± standard deviation (SD) from three independent experiments performed in triplicate.

| Sl. No. | Cancer type | Cell line | Gloriosine, IC<sub>50</sub> (nM), Stdv. |
| --- | --- | --- | --- |
| 1 | Breast cancer | MDA-MB-231 | 100.65 ± 8.29 |
| 2 | Breast cancer | Hs 578T | 99.82 ± 7.61 |
| 3 | Breast cancer | HCC1395 | 88 ± 5.64 |
| 4 | Normal breast cells | MCF 10A | 720.83 ± 18.19 |
| 5 | Gastric cancer | KATO III | 141.52 ± 8.89 |
| 6 | Gastric cancer | AGS | 109.20 ± 7.8 |
| 7 | Ovarian cancer | SK-OV-3 | 120.91 ± 18.19 |
| 8 | Ovarian cancer | Caov-3 | 101.53 ± 10.52 |

### Gloriosine inhibits glutamine utilization and induces ferroptosis cell death in TNBC

To elucidate the mechanism underlying gloriosine-mediated cell death, the involvement of glutamine metabolism was first examined. Western blot analysis revealed a concentration-dependent downregulation of GLS1 (Figure 7A), a key enzyme in glutaminolysis, responsible for the conversion of glutamine to glutamate, following gloriosine treatment. Furthermore, determination of the glutamine/glutamate ratio in the gloriosine-treated sample suggested potentially impaired glutathione biosynthesis and contributed to redox imbalance in both cells (Figure 7B). Subsequently, intracellular ROS levels were assessed using DCFDA staining. Treatment with gloriosine induced a robust, concentration-dependent increase in ROS generation in both cell types (Figure 7C). Consistent with these observations, intracellular GSH levels were significantly depleted following gloriosine treatment (Figure 7D), indicating disruption of cellular redox homeostasis. Moreover, MDA analysis study further supported the lipid peroxidation in cells (Figure 7E).

**Figure 7:**
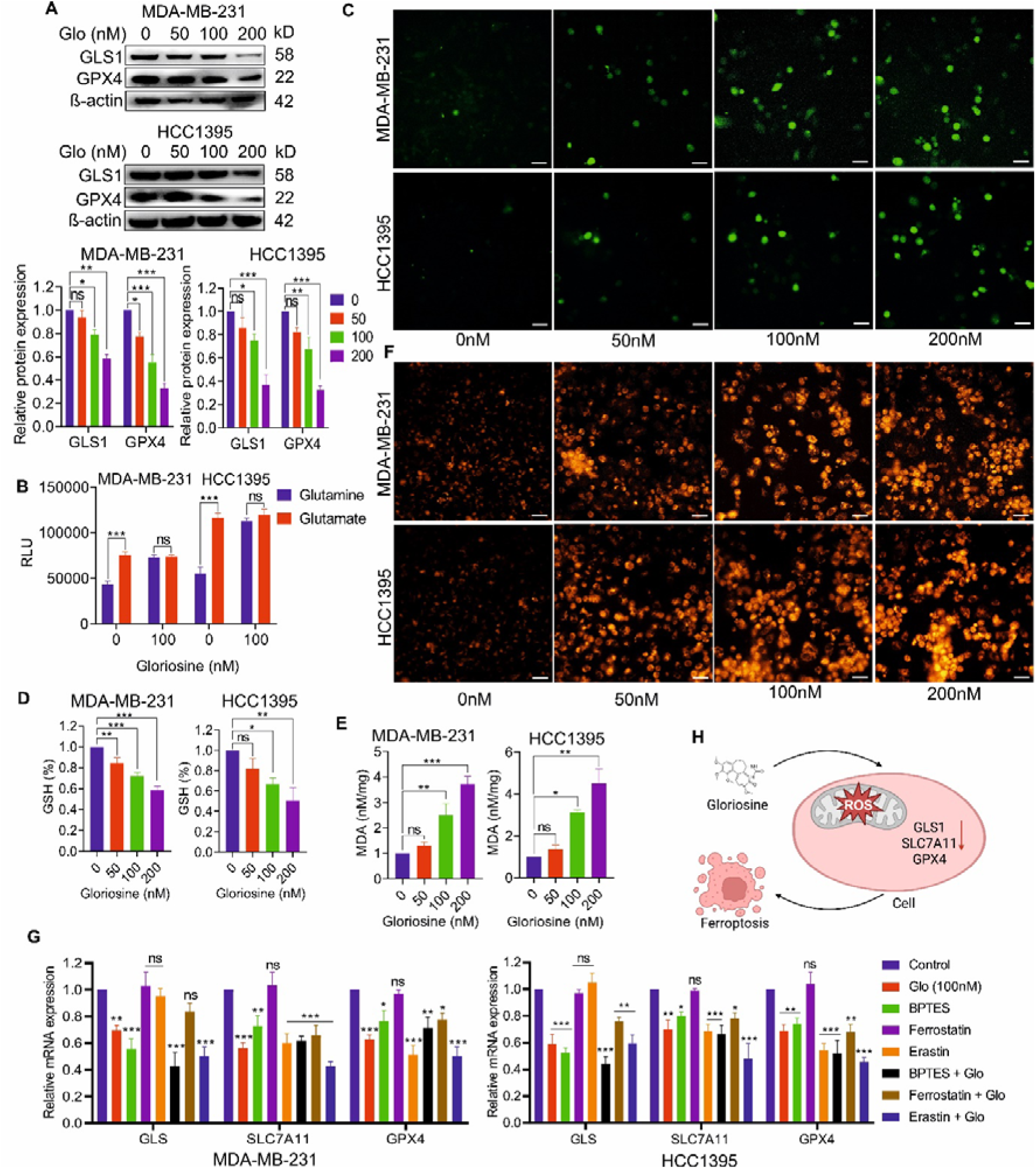
Gloriosine induces ferroptotic cell death in MDA-MB-231 cells. **(A)** Western blot analysis showing the expression levels of GLS1 and GPX4 following treatment with increasing concentrations of gloriosine; β-actin served as a loading control. **(B)** MDA-MB-231 and HCC1395 cells were collected and lysed as described in the Materials and Methods section. The cell lysates were titrated and then used to measure metabolites in three replicate wells. The mean relative light units (RLU) are shown for glutamine and glutamate detection using 4000 cells/well. **(C)** Representative fluorescence microscopy images of DCFDA-stained cells after 48 h of gloriosine treatment (scale bar = 25 µm), indicating intracellular ROS accumulation. **(D)** Quantification of reduced glutathione (GSH) levels in cells treated with varying concentrations of gloriosine. **(E)** Quantification of malondialdehyde (MDA) levels under the same treatment conditions. **(F)** FerroOrange staining demonstrating intracellular ferrous iron accumulation in MDA-MB-231 cells after 48 h of treatment (scale bar = 25 µm). Data are presented as mean ± SD from three independent experiments. Statistical significance was determined using one-way ANOVA (*P < 0.05, **P < 0.01, ***P < 0.001). **(G)** mRNA expression levels of SLC7A11 and GPX4 were analyzed in breast cancer cells following treatment with gloriosine alone or in combination with BPTES, ferrostatin-1, or erastin. Gene expression was quantified by qRT-PCR and normalized to GAPDH. **(H)** Overall summary represented in Fig. 7. Abbreviations: Glo: gloriosine, ferrostatin: ferrostatin 1.

To investigate whether this effect was associated with ferroptosis, Western blot analysis was performed, which revealed a gradual, concentration-dependent reduction in GPX4 expression, a key regulator of ferroptosis (Figure 7A). Additionally, FerroOrange staining data further supported these findings, confirming enhanced ferroptotic activity in TNBC cells upon gloriosine treatment (Figure 7F).

For further confirmation of the role of ferroptosis in gloriosine-induced cell death, breast cancer cells were treated with gloriosine alone or in combination with erastin, or ferrostatin-1. Notably, co-treatment with erastin potentiated the cytotoxic effects of gloriosine, whereas ferrostatin-1 significantly rescued cells from gloriosine-induced death (Figure 7G). These results demonstrated that induction of ferroptosis enhances gloriosine sensitivity, while pharmacological blockade of ferroptosis mitigates its effects, supporting a ferroptosis-dependent mechanism of gloriosine activity. A schematic summary of the proposed mechanism is presented in Figure 7H.

## Discussion

Conventional chemotherapy remains the primary treatment option for TNBC; however, its clinical efficacy is often limited by systemic toxicity, drug resistance, and high rates of recurrence and metastasis [1][2]. These challenges underscore the urgent need for the development of novel chemotherapeutic agents that are more selective, less toxic, and capable of targeting alternative cell death pathways. Natural compounds have emerged as promising candidates in this context, owing to their multi-targeted mechanisms and relatively lower toxicity profiles. In this regard, we have isolated gloriosine, a bioactive alkaloid, which showed antiproliferative properties against different cancer cells with minimal toxicity to normal breast cells (Table 1) [32], suggesting its potential as a promising therapeutic candidate.

This study combines network pharmacology and experimental validation to explore the therapeutic potential and mechanisms of gloriosine in TNBC. By combining in-silico target prediction with biological assays, our findings provide a comprehensive understanding of how gloriosine exerts its anti-cancer effects, particularly in the context of glutamine metabolism and ferroptosis.

Target prediction analysis identified 100 potential protein targets of gloriosine, which were further refined through standardization, ensuring a reliable dataset for downstream analysis. Intersection of these targets with TNBC-associated and glutamine metabolism-related genes resulted in 60 overlapping targets, suggesting that gloriosine may exert its therapeutic effects by modulating key molecular nodes that link cancer progression with metabolic reprogramming. Given the well-established role of glutamine metabolism in supporting rapid proliferation and survival of TNBC cells [33][34], these findings suggest the potential of gloriosine as a metabolic modulator.

The PPI network further emphasized the importance of several hub genes, particularly SRC, EGFR, mTOR, and HSP90AA1, which exhibited the highest degree values. Their high connectivity suggests that gloriosine may exert multi-target effects by influencing critical signaling hubs, thereby disrupting the complex regulatory networks that sustain TNBC progression. Aberrant activation of SRC signaling is frequently observed in TNBC and is associated with enhanced tumor aggressiveness and poor clinical outcomes [35]. SRC interacts with receptor tyrosine kinases such as EGFR, amplifying downstream PI3K/AKT and MAPK signaling pathways that promote tumor growth and metastasis [36]. In addition to its role in tumor progression, SRC also contributes to metabolic reprogramming, including the regulation of glutamine metabolism to support anabolic growth and redox balance in cancer cells [37][38]. EGFR is commonly overexpressed in TNBC and is linked to poor prognosis, increased proliferation, and metastatic potential. Its activation drives key downstream signaling pathways, including PI3K/AKT and MAPK [39][40], and enhances glutamine uptake and utilization by regulating transporters and metabolic enzymes, thereby supporting rapid tumor growth [41][42]. Similarly, aberrant activation of the PI3K/AKT/mTOR pathway plays a central role in TNBC progression and therapeutic resistance [43]. mTOR, in particular, is a key regulator of glutamine metabolism, promoting its uptake and utilization to sustain cellular biosynthesis and redox homeostasis [44]. Additionally, elevated expression of HSP90AA1 is associated with poor prognosis and tumor aggressiveness in TNBC, as it stabilizes multiple oncogenic proteins and supports critical signaling pathways such as PI3K/AKT, MAPK, and ERK [45][46][47]. The presence of moderately connected targets such as FPR1, CYP19A1, and MAPK9 further supports the idea that gloriosine impacts diverse biological processes.

GO enrichment analysis revealed that the identified targets are primarily involved in metabolic processes, biological regulation, and responses to stimuli, reinforcing the hypothesis that gloriosine interferes with cancer-associated metabolic and regulatory pathways. Cellular component analysis indicated localization in key cellular compartments such as the nucleus and membrane systems, while molecular function analysis highlighted roles in protein binding and catalytic activities. These findings collectively suggest that gloriosine may influence multiple layers of cellular function, contributing to its anti-cancer activity.

KEGG pathway enrichment analysis identified several critical pathways, including EGFR tyrosine kinase inhibitor resistance, central carbon metabolism, prolactin signaling, ErbB signaling, and endocrine resistance. These pathways are closely associated with TNBC aggressiveness, therapeutic resistance, and metabolic adaptation [39][48][49][50]. The involvement of these pathways suggests that gloriosine may not only inhibit tumor growth but also potentially overcome resistance mechanisms, which is a major challenge in TNBC treatment.

Based on the degree value form PPI network, molecular docking studies were performed to further validate the interaction between gloriosine and key hub targets, including SRC, EGFR, mTOR, and HSP90AA1. The docking results demonstrated that gloriosine exhibited strong binding affinities toward these proteins, forming stable interactions within their active binding sites, with the highest binding compatibility observed for HSP90AA1. These findings suggest that gloriosine can effectively bind to and potentially modulate the activity of these critical signaling proteins, thereby supporting its multi-target therapeutic potential in TNBC.

Experimental validation further substantiated the computational predictions. Gloriosine demonstrated potent cytotoxicity in TNBC cell lines (MDA-MB-231, HCC1395 and Hs 578T), with significantly lower IC□□ values compared to normal breast epithelial cells, indicating selective anti-cancer activity. The observed concentration-dependent reduction in cell viability, cell proliferation and inhibition of migration confirm its anti-proliferative and anti-metastatic potential.

In addition to its cytotoxic effects, gloriosine appears to interfere with glutamine metabolism at the cellular level. Specifically, our findings suggest that gloriosine inhibits GLS1, thereby limiting glutamine utilization and restricting the supply of critical metabolic intermediates required for anabolic growth and redox balance in TNBC cells. Moreover, our study provides strong evidence that gloriosine induces ferroptosis in TNBC cells. The marked increase in intracellular reactive oxygen species (ROS) and MDA levels, coupled with depletion of glutathione (GSH), indicates disruption of redox homeostasis and lipid peroxidation, a hallmark of ferroptotic cell death. Furthermore, the downregulation of GPX4, a key enzyme that protects cells from lipid peroxidation, strongly supports the induction of ferroptosis. The FerroOrange staining results, indicating increased intracellular iron levels, further validate this mechanism. In addition, our study showed that co-treatment with the ferroptosis inducer erastin significantly potentiated the anticancer effects of gloriosine, whereas the ferroptosis inhibitor ferrostatin-1 markedly attenuated its activity. The ability of ferrostatin-1 to rescue cells from gloriosine-induced cytotoxicity strongly supports the notion that ferroptosis is a major mechanism underlying its anticancer effects.

Despite providing significant insights, this study has several limitations. First, the target identification and pathway analyses were primarily based on in silico approaches, which may be subject to database bias and prediction inaccuracies. Second, although *in vitro* experiments validated the anti-cancer effects of gloriosine, these findings are limited to cell line models and should be validated on an *in vivo* model. Third, although our findings suggest that gloriosine induces ferroptosis, the precise molecular signaling underlying these effects remains unexplored. Additionally, the molecular docking results indicate potential interactions between gloriosine and key targets; however, these interactions were not further confirmed using biophysical or structural validation techniques.

## Conclusion

In conclusion, the present study demonstrates that gloriosine possesses significant anti-cancer potential against TNBC through a multi-target and multi-pathway mechanism. By integrating network pharmacology, molecular docking, and experimental validation, key targets and signaling pathways involved in the therapeutic action of gloriosine were identified, particularly those associated with cancer metabolism and drug resistance. The findings indicate that gloriosine effectively inhibits TNBC cell proliferation and migration, along with the induction of ferroptotic cell death. Collectively, this study provides a comprehensive framework suggesting that gloriosine may serve as a promising therapeutic candidate for TNBC by targeting critical oncogenic pathways and promoting ferroptosis.

## Acknowledgement

Authors are grateful to the Council of Scientific and Industrial Research (CSIR), New Delhi, India, for supporting the research fellowship. Authors are also thankful to the National Institute of Pharmaceutical Education and Research Hyderabad (NIPER Hyderabad), Telangana, India for instrumental facility. The authors also gratefully acknowledge Chandrima Saha for her valuable assistance with the network pharmacology analysis.

## Authors contributions

B.D. was involved in conceptualization, data curation, formal analysis, methodology, validation, and writing – original draft. B.D. and S.K.G. were involved in project administration. E.C. reviewed the manuscript. B.G. synthesized the compound and performed the docking study. S.K.J. supervised the work. P.K.N. and S.K.G. were involved in project Investigation. S.K.G. was involved in funding acquisition, investigation, project administration, providing resources, visualization, and reviewing and editing the manuscript. All authors reviewed the manuscript at its final form.

## Data availability statement

The data that support the findings of this study are available from the corresponding author upon request.

## Abbreviations

ABCC1: ATP Binding Cassette Subfamily C Member 1
ADAM17: ADAM Metallopeptidase Domain 17
CAPN1: Calpain 1
CTSL: Cathepsin L
CYP19A1: Cytochrome P450 Family 19 Subfamily A Member 1
EGFR: Epidermal Growth Factor Receptor
ErbB: Erythroblastic Leukemia Viral Oncogene Homolog
FLT4: Fms Related Receptor Tyrosine Kinase 4
FPR1: Formyl Peptide Receptor 1
GLS1: Glutaminase 1
GPX4: Glutathione Peroxidase 4
GRM1: Glutamate Metabotropic Receptor 1
HSP90AA1: Heat Shock Protein 90 Alpha Family Class A Member 1
LRRK2: Leucine-Rich Repeat Kinase 2
MAPK9: Mitogen-Activated Protein Kinase 9
MMP1: Matrix Metallopeptidase 1
mTOR: Mechanistic Target of Rapamycin
NOS2: Nitric Oxide Synthase 2
NR3C2: Nuclear Receptor Subfamily 3 Group C Member 2
P2RX7: Purinergic Receptor P2X 7
PARP3: Poly(ADP-ribose) Polymerase 3
PDE4D: Phosphodiesterase 4D
PGK1: Phosphoglycerate Kinase 1
SRC: Non-Receptor Tyrosine Kinase
TGM2: Transglutaminase 2

